# The Limits of Cross-Species WGCNA: Library Imbalance and Signal Dilution Constrain Effector Gene Recovery in Dual-Organism RNA-seq

**DOI:** 10.64898/2026.04.30.721941

**Authors:** Amit Fenn, Ralph Hückelhoven, Nadia Kamal

## Abstract

Dual-organism RNA sequencing (RNA-seq) experiments, in which the transcriptomes of a host and a microbe are sequenced simultaneously, are increasingly used to study plant–microbe interactions. A central analytical goal is identifying effector proteins and their host targets through gene co-expression. Weighted Gene Co-expression Network Analysis (WGCNA) is the dominant tool for gene co-expression analyses, yet its ability to recover interaction-interface genes from a merged dual-organism matrix has not been systematically characterised. Here we present a simulation framework using real gene models from *Hordeum vulgare* (barley) and *Blumeria graminis* f. sp. *Hordei* M.Liu & Hambl (powdery mildew) to evaluate single-network WGCNA across a gradient of plant-to-fungal library size ratios (1:1–20:1), three levels of co-expression signal strength, and three WGCNA network construction types (signed, unsigned, signed hybrid). We embed 20 model effector genes (bridge genes) driven by a mixed host–pathogen eigengene and evaluate recovery using four metrics aligned with the biological objective: cross-species hub rank, top-decile hub enrichment, bridge gene detection rate, and bridge co-separation (the fraction of effector–target pairs co-assigned to the same detected module). Across 225 simulation runs (15 conditions × 5 replicates × 3 network types), bridge genes are robustly identifiable as cross-species connectivity hubs (mean rank 0.92 versus 0.50 for module genes) but co-assignment of effector–target pairs to the same module fails in 41% of runs due to scale-free topology collapse. Signal strength (*η*^2^ = 0.12) and library ratio (*η*^2^ = 0.22) are the primary determinants of co-separation, while network type choice accounts for less than 2%. A read-depth bias systematically inflates pathogen gene hub ranks relative to host genes at high ratios. These results establish that the method can identify effector candidates as cross-species hubs under a broad range of conditions, but reliable co-assignment requires adequate pathogen read depth and strong co-expression signal—properties that experimental design, not analytical parameterisation, must provide.

## 1 Introduction

Studies that profile transcriptomes from only one partner in a plant–microbe interaction cannot capture the reciprocal gene expression dynamics that unfold on both sides simultaneously. Dual species RNA-seq addresses this limitation by sequencing host and pathogen RNA from the same biological sample, producing a single count matrix in which both transcriptomes are observed under identical experimental conditions [Westermann et al., 2012]. The approach has been applied across diverse plant–pathogen systems: Hacquard et al. simultaneously profiled *Blumeria graminis* f.sp. *hordei* and its hosts barley and immunocompromised arabidopsis across four early infection stages, revealing a conserved fungal transcriptional programme despite the evolutionary distance between hosts [Hacquard et al., 2013], while Nobori et al. used a similar strategy in the *Pseudomonas syringae*–*Arabidopsis* system to map the bacterial transcriptome across 27 plant genotype and bacterial strain combinations [Nobori et al., 2018]. In each case, reads were mapped independently to each species’ reference genome and analysed using differential expression within species. The resulting data contain expression measurements for thousands of host and pathogen genes across the same set of samples—conditions that naturally invite the question of whether co-expression structure can be detected across both organisms jointly, and in particular whether effector proteins and their host targets leave a detectable signal of transcriptional co-regulation.

Effector proteins are secreted by pathogens to manipulate host cell processes and suppress immunity [Bozkurt et al., 2012]. Their discovery has historically relied on genome-wide secretome prediction followed by functional screening for immune suppression activity. Transcriptional co-expression between secreted effector candidates and the host genes they target offers a complementary detection strategy: an effector deployed during infection to modulate a specific host pathway should be co-expressed—across infection stages and conditions—with the genes in that pathway. If the dual RNA-seq count matrix can be used to identify genes that are simultaneously strongly co-expressed with both the pathogen transcriptome and the host immune response, such genes could be strong effector candidates without requiring prior knowledge of secretion signals or functional assays.

Weighted gene co-expression network analysis (WGCNA) [Langfelder and Horvath, 2008] is the most widely used framework for identifying modules of co-expressed genes. WGCNA offers three network construction types—*signed, unsigned*, and *signed hybrid* —which differ in how they transform correlation coefficients into adjacency weights. The method was developed and benchmarked on single-organism expression data, and the developers note that results become unreliable in the presence of mixed-source inputs [Langfelder and Horvath, 2008]—conditions that precisely describe a merged dual-organism matrix. Benchmarking of co-expression network construction from RNA-seq data has been conducted systematically, but exclusively in single-organism settings [Johnson and Krishnan, 2022]. Whether joint single-network WGCNA can detect interaction-interface genes in dual-organism data, and what conditions limit this, has not been evaluated.

Applying WGCNA to a merged dual-organism matrixraises a question that has not previously been addressed: can the method identify genes whose co-expression spans both organisms, and specifically can it recover the transcriptional signal of an effector protein that is simultaneously co-regulated with pathogen infection programmes and host immune responses? This is a non-trivial question for several reasons. The two organisms contribute gene sets of different sizes and expression variance; sequencing depth is split between genomes in proportions that vary with infection stage and host resistance; and the scale-free topology criterion that guides soft-threshold selection was developed for single-organism networks, where all genes belong to the same co-expression landscape. Whether these properties of the merged matrix preserve or destroy the cross-species co-expression signal of interaction-interface genes, and what experimental conditions govern this, is precisely what a simulation benchmark can establish. A module containing both a barley immune receptor and the *Blumeria* effector suppressing it is not a methodological artefact to be explained away: it is the target output of an interaction-aware analysis, and determining when WGCNA reliably produces such modules is the central question of this work.

Here, we address this question with a systematic simulation study grounded in the biology of barley powdery mildew (*B. graminis* f.sp. *hordei*). We generate synthetic count matrices for barley (*H. vulgare*) and *Blumeria hordeii* using the respective species-specific gene models gene models and realistic co-expression module structure, embedding 20 model effector genes (bridge genes) driven by a mixed eigengene with equal loading on both organisms’ transcriptional programmes. We vary the library size ratio between the two organisms across five levels (1:1 to 20:1), vary co-expression signal strength across three levels, and evaluate the ability of joint single-network WGCNA—using all three standard network types—to recover these model effectors as cross-species connectivity hubs and to co-assign them with their interaction partners into the same detected modules. Our simulation is evaluated against four metrics aligned with the biological objective: cross-species hub rank, top-decile hub enrichment, bridge gene detection rate, and bridge co-separation (the fraction of effector– target pairs co-assigned to the same detected module).

We find that bridge genes are robustly identifiable as cross-species connectivity hubs across the full range of tested conditions—mean hub rank 0.92 versus 0.50 for ordinary module genes—but that their co-assignment into shared modules is fragile, failing in 41% of runs due to network collapse driven by the scale-free topology constraint. Signal strength and library ratio jointly determine whether co-assignment succeeds (*η*^2^ = 0.12 and 0.22 respectively), while network type choice has minimal impact on co-separation (*η*^2^ *<* 0.02) despite strongly influencing hub rank (*η*^2^ = 0.18). A critical asymmetry emerges at the gene level: pathogen bridge genes achieve higher cross-species hub rank than host bridge genes across all tested ratios, not because of biological asymmetry but because the ranking is biased by read depth—pathogen genes are ranked by their connections to the deep host side of the matrix. Together these results reframe the limits of joint WGCNA in dual-organism settings: the method can identify effector candidates as cross-species hubs under a broad range of conditions, but reliable co-assignment with host targets requires adequate pathogen read depth and strong co-expression signal— properties that experimental design, not analytical parameterisation, must provide.

## 2

## 3 Results

### 3.1 Simulation design overview

The simulation generated paired barley-*Blumeria* expression datasets with planted ground-truth co-expression structure across a factorial design of library ratio and signal strength. Each dataset comprised 60 samples drawn from five infection timepoints with twelve biological replicates per timepoint, with module eigengene trajectories following an AR(1) process (ϕ=0.7) at the timepoint level and replicate-level noise (σ=0.3) added within each timepoint (Fig.1A). The gene pool consisted of 625 barley genes distributed across five modules and a grey background, and 375 *Blumeria* genes across three modules and background, with 20 bridge genes embedded within the module gene pool as model effectors driven by a mixed host--pathogen eigengene (Fig.1B). Library ratio was implemented by partitioning a fixed total sequencing depth of ~10M reads per sample between species, ranging from 1:1 to 20:1 (Fig.1C). Across the full factorial design, 75 independent expression datasets were generated spanning five ratios and three signal levels, each analysed under all three WGCNA network types for a total of 225 runs (Fig.1D).

**Figure 1:**
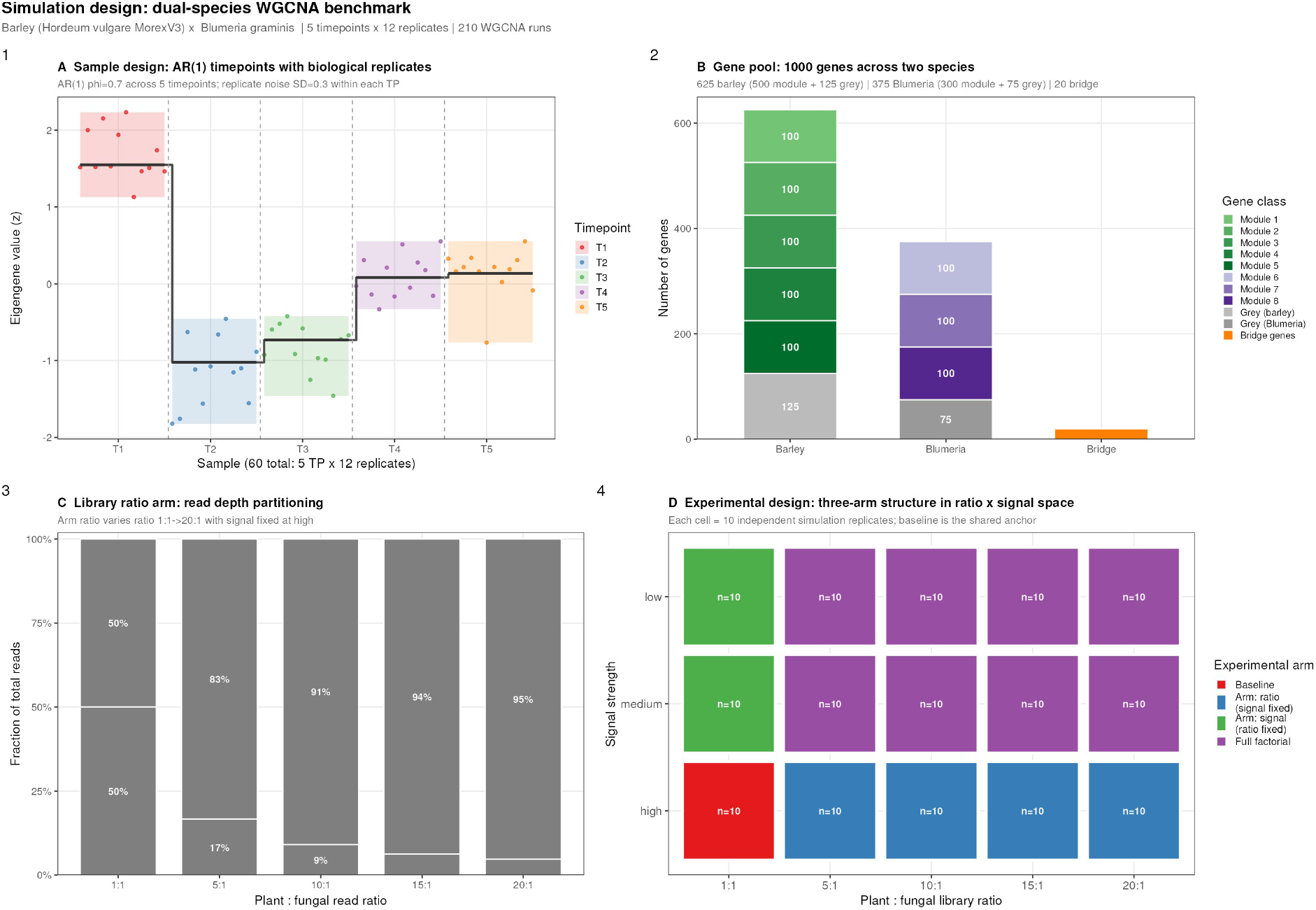
Simulation design overview. **(A)** Sample structure: 5 timepoints × 12 replicates. Module eigengenes follow AR(1) trajectories (*ϕ* = 0.7) at the timepoint level; replicate noise (SD = 0.3) is added within each timepoint. **(B)** Gene pool: 1,000 total genes split into barley (625) and *Blumeria* (375) transcriptomes, with 20% grey genes per species and 20 bridge genes (model effectors). **(C)** Library ratio arm: total sequencing depth held constant; plant-to-fungal read allocation varies from 1:1 to 20:1. **(D)** Experimental design: four-arm structure (baseline, ratio arm, signal arm, full factorial) spanning 5 ratios × 3 signal levels, each replicated 5 times and analysed under 3 network types.

### 3.2 Bridge genes as model effectors

To simulate the transcriptional behaviour of effector proteins at the host-pathogen interface, we embedded 20 bridge genes into the gene pool 10 barley and 10 *Blumeria* whose expression was driven by a mixed eigengene constructed as an equal-weighted combination of the first barley and first *Blumeria* module eigengenes. This gives bridge genes moderate co-expression with both species simultaneously: they correlate with the infection programme on the fungal side and with the immune response on the host side, without belonging exclusively to either. This is the transcriptional signature we would expect from a secreted effector that is upregulated during active infection and targets a specific host pathway, co-regulated with the pathogen’s deployment programme and with the host genes it manipulates. Bridge genes were drawn from within the existing module gene pool and overwritten at the expression construction stage, so the total gene pool size is unchanged and bridge genes compete for module assignment on equal terms with all other genes. Their ground-truth class is distinct from both module genes (pure single-species co-expression) and grey genes (no co-expression structure)

### 3.3 Bridge genes are recoverable as cross-species hubs

We first asked whether bridge genes—model effectors driven by a mixed eigengene with equal loading on both species—emerge as identifiable hubs in the cross-species connectivity landscape of the merged expression matrix (Fig. 2).. Bridge genes consistently occupy the right tail of the distribution, achieving a mean rank of 0.92 across all conditions versus 0.50 for module genes and 0.45 for grey genes. At the best-case condition (ratio 1:1, high signal), 76% of bridge genes fall in the top decile of cross-species connectivity—7.6-fold above the 10% random expectation. This hub enrichment is preserved across all five ratio levels: mean rank remains between 0.87 and 0.94 regardless of library imbalance, because as the ratio increases, fungal module genes lose read depth and their within-species correlations compress, narrowing the competition for the top of the *k*_cross_ ranking and pushing bridge genes—which retain their connections to the abundant barley side—upward in relative rank.

**Figure 2:**
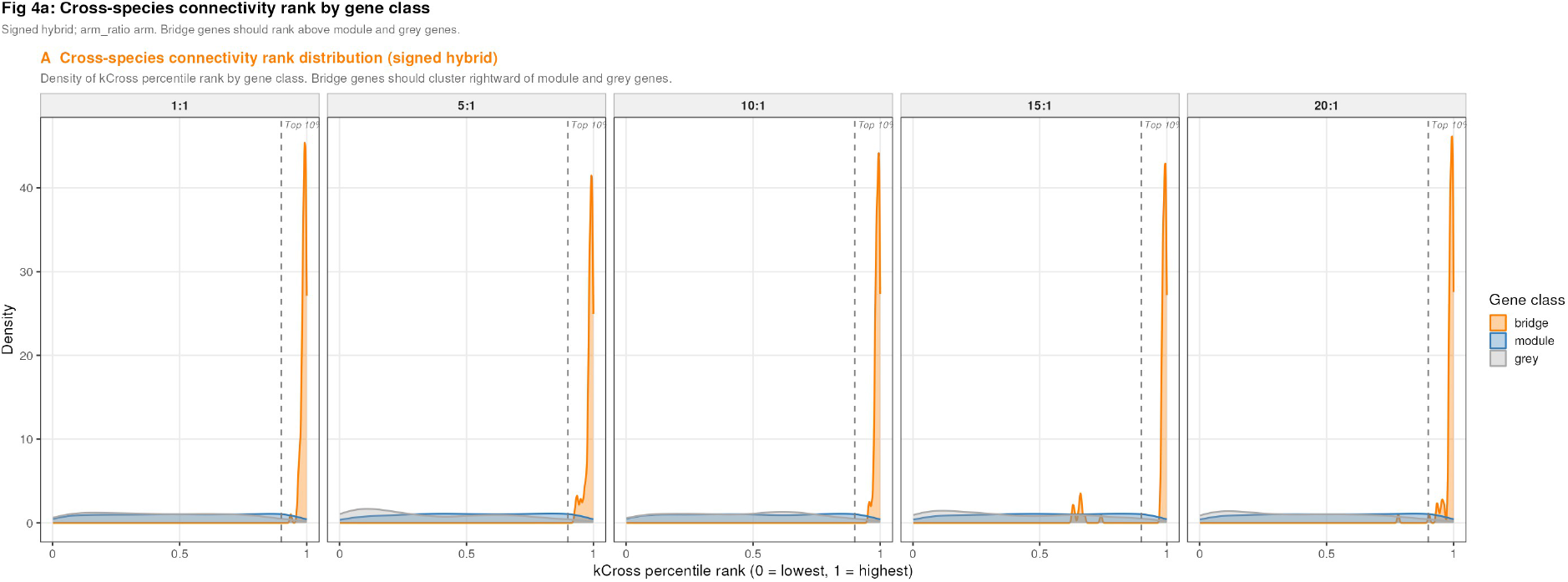
Cross-species connectivity rank of bridge genes. Density of *k*_cross_ percentile rank for bridge (orange), module (blue), and grey (grey) genes under signed hybrid at representative library ratios. The dashed vertical line marks the 90th percentile. Bridge genes consistently occupy the right tail regardless of ratio, while module and grey genes cluster near the centre of the distribution.

### 3.4 Module collapse concentrates at low biological signal and is detectable prospectively

### 3.5

In 92 of 225 simulation runs (41%), WGCNA detected one module or none, making co-assignment of effector target pairs impossible regardless of hub status. This collapse is driven by the soft-power selection procedure: when neither species’ correlation landscape satisfies the scale-free topology criterion over the tested power range, the fallback to the highest-R2 power produces an extreme adjacency transformation that compresses most genes into a single dense clique or background. Crucially, collapse is not uniformly distributed: it is concentrated at low co-expression signal (47--93% collapse rate across ratios at low signal, Fig.~6) and is largely independent of network type (Figs.~S2 and S3). This means collapse is a diagnostic about the underlying biological signal, not a random failure of the algorithm, and can be detected prospectively by inspecting the soft-power curve and module count before results are interpreted. Paradoxically, bridge gene kcross rank is *higher* in collapsed runs (mean 0.96) than in successful runs (0.89), because the compressed adjacency matrix retains genuine cross-species signal preferentially while suppressing within-species module structure. This has a practical consequence: hub ranking remains an informative effector candidacy screen even in collapsed experiments where co-assignment is unavailable, but the hub rank should not be taken as evidence that the network has successfully resolved co-expression structure.

### 3.6 BBridge gene co-separation quantifies effector--target co-clustering and defines the operating envelope of the method

Bridge co-separation — the fraction of cross-species bridge gene pairs (10 plant × 10 fungal = 100 pairs) assigned to the same detected non-grey module — is the metric most directly aligned with the biological objective: it measures whether model effectors and their host interaction partners are placed in the same co-expression community (Fig.~5). At the best-case condition (ratio 1:1, high signal), mean co-separation is 0.72, meaning nearly three-quarters of effector--target pairs are correctly co-clustered — a result that directly supports functional hypothesis generation from dual-organism co-expression data. Co-separation remains above 0.50 up to a 10:1 library ratio at high signal, covering the majority of published barley-*Blumeria* infection datasets. At more challenging conditions — ratio 20:1 or low co-expression signal — co-separation falls to 0.32 and 0.19 respectively, indicating that under these circumstances hub ranking alone (Section~3.2) is the more reliable effector candidacy screen. Taken together, the co-separation results define a practical operating envelope: experiments with manageable library ratio and sufficient biological signal yield co-expression communities that directly implicate effector--target pairs, while experiments outside this envelope still yield hub-based candidates through the kcross ranking tier.

**Figure 3:**
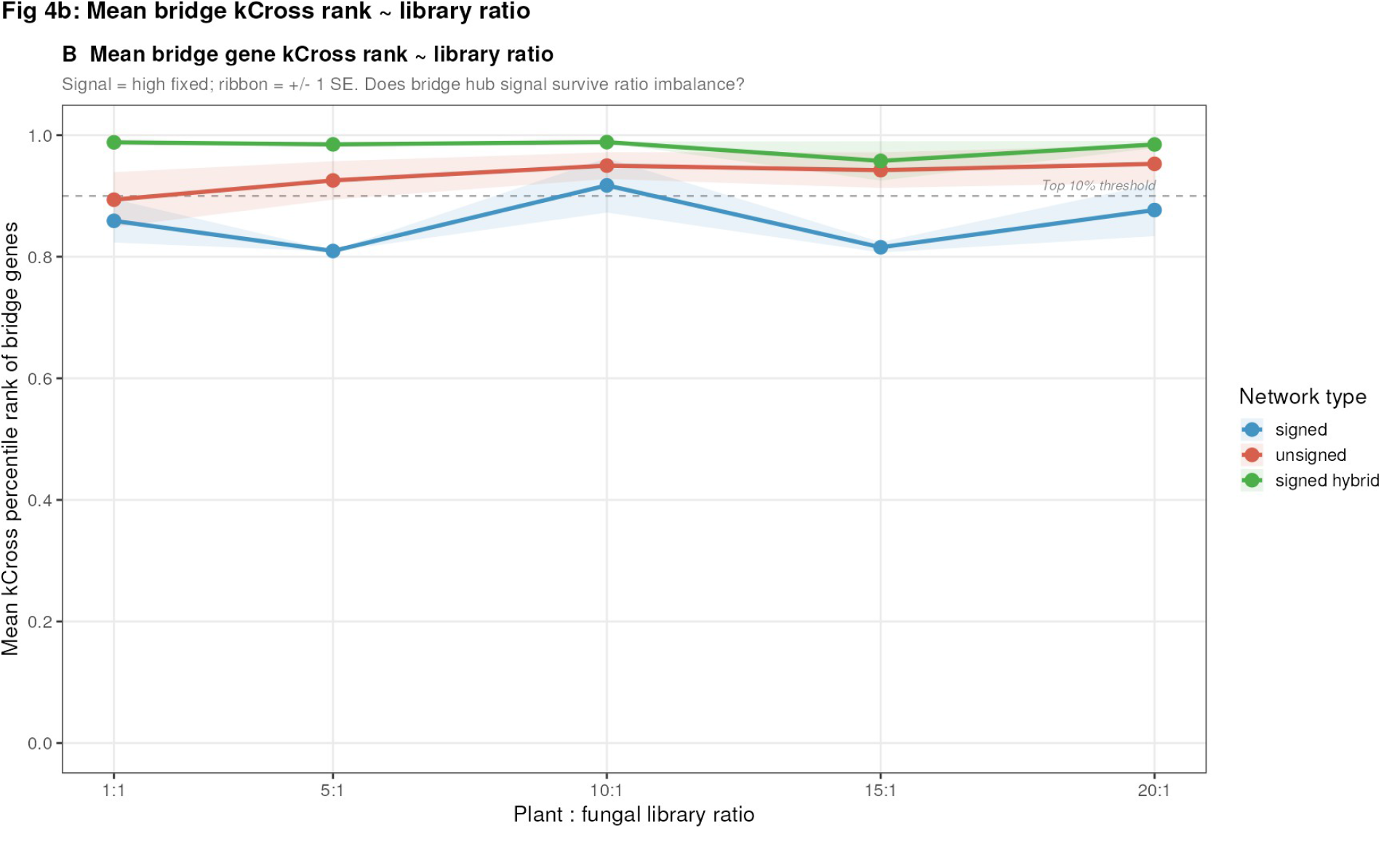
Mean bridge gene *k*_cross_ rank across library ratios. Mean percentile rank of bridge genes in the cross-species connectivity distribution for each network type (signal = high fixed). Ribbon = *±*1 SE. The dashed line marks the 90th-percentile threshold. Hub rank is preserved across ratios for all three network types, with signed hybrid consistently highest.

**Figure 4:**
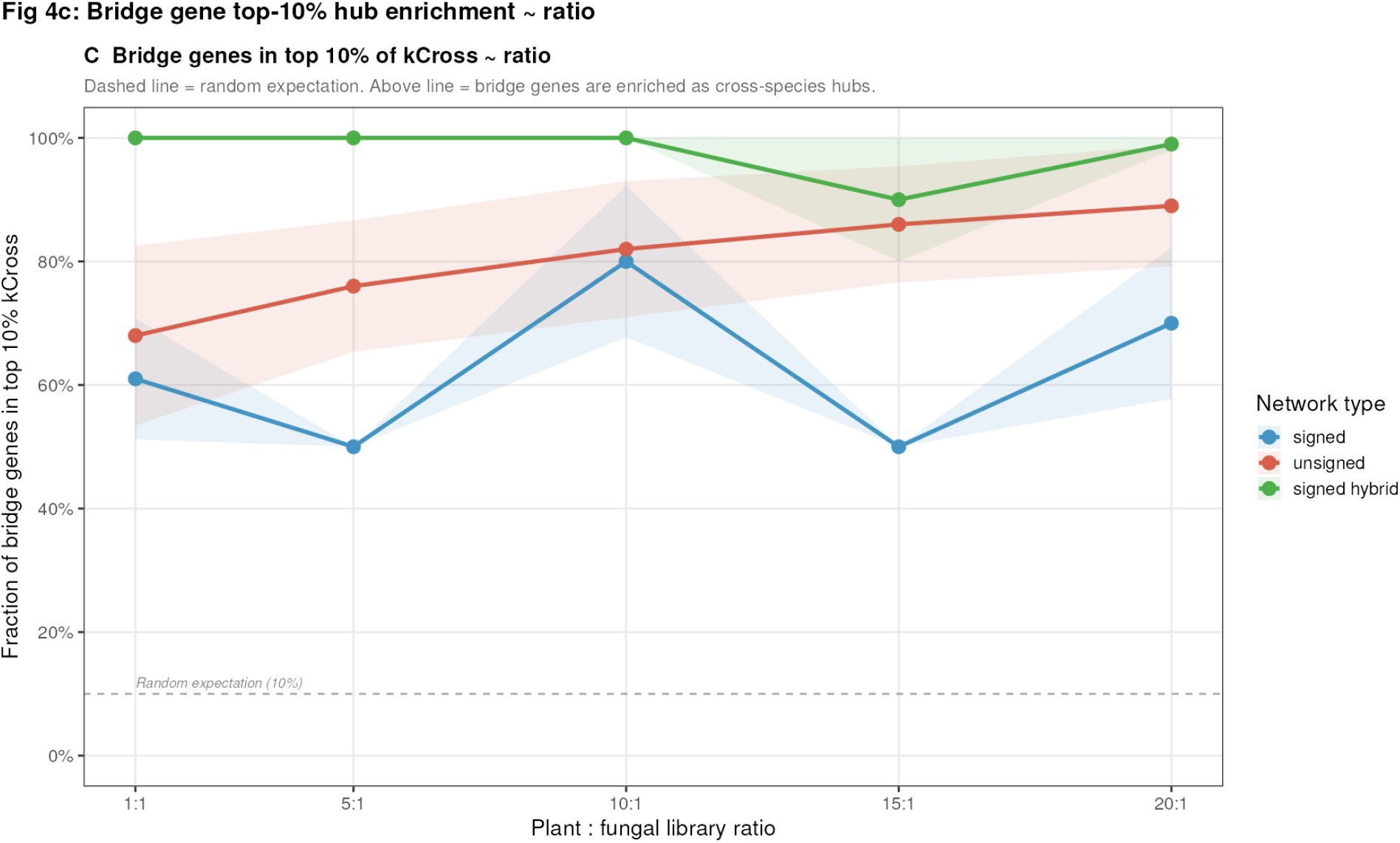
Bridge gene enrichment in top 10% of cross-species hubs. Fraction of bridge genes in the 90th percentile or above of *k*_cross_, by library ratio and network type (signal = high fixed). The dashed line marks the random expectation (10%). All network types show substantial hub enrichment across the full ratio range.

**Figure 5:**
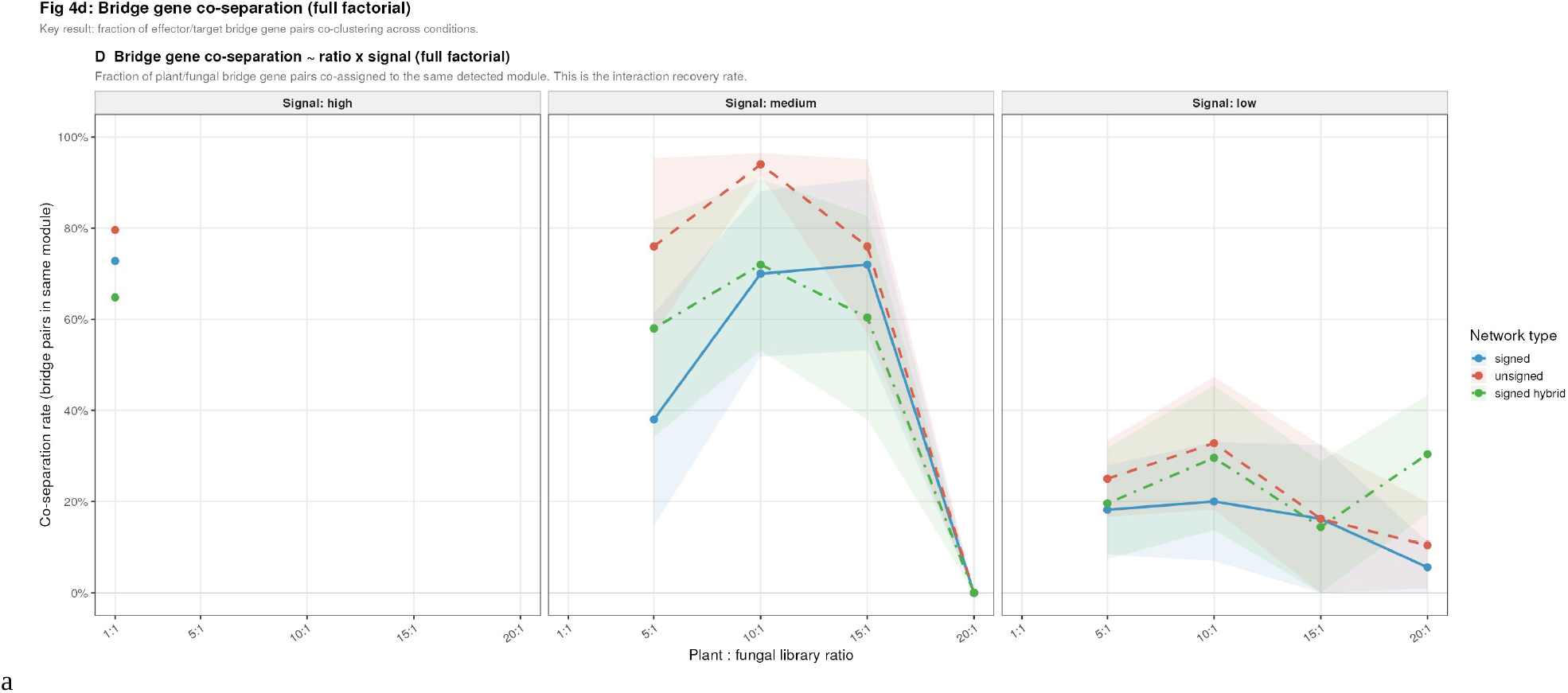
Bridge gene co-separation across ratio × signal space. Fraction of crossspecies bridge gene pairs (model effector + target) assigned to the same detected non-grey module, shown by library ratio for each signal level (panels) and network type (colour). Co-separation is highest at low ratio and high signal; all three network types produce comparable profiles.

### 3.7 Signal strength and library ratio are the dominant determinants of bridge gene recovery

The variance decomposition (Fig. 6C) shows that library ratio and signal strength jointly account for the majority of variance in primary recovery metrics. For bridge co-separation, ratio explains 22% of total variance (*η*^2^ = 0.22), signal strength 12% (*η*^2^ = 0.12), and their interaction a further 10% (*η*^2^ = 0.10). Network type explains less than 2% of variance in co-separation (*η*^2^ = 0.011). The ratio × signal interaction reflects a non-additive effect: at low signal, increasing the library ratio has essentially no further impact because the method has already collapsed (Fig. 6A–B)

**Figure 6:**
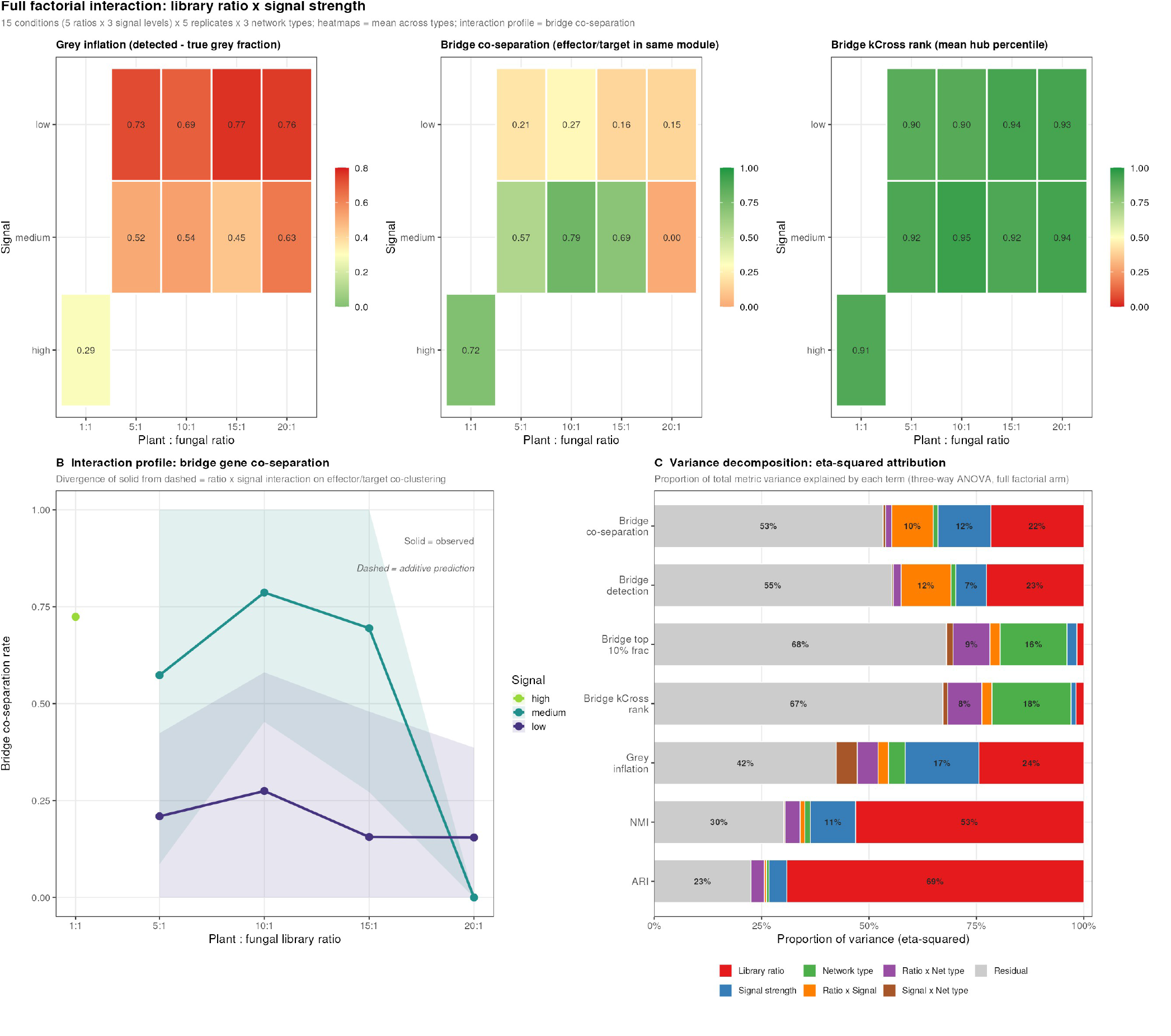
Full factorial interaction and variance decomposition. **(A)** Grey inflation across ratio × signal space, averaged across network types. **(B)** Bridge co-separation as an interaction profile; solid lines = observed, dashed = additive prediction. Divergence between solid and dashed confirms a ratio × signal interaction: low signal damages co-separation independently of ratio. **(C)** Eta-squared (*η*^2^) variance decomposition from three-way ANOVA. Ratio and signal dominate all bridge recovery metrics; network type accounts for *<*2% of variance in co-separation and detection rate.

### 3.8 One-at-a-time arm analysis confirms ratio and signal as independent limiting factors

### 3.9 Network type strongly influences hub rank but not co-assignment

Although network type has minimal impact on co-separation, it has a pronounced effect on bridge hub rank: *η*^2^ = 0.18 for *k*_cross_ rank, larger than the effects of ratio (*η*^2^ = 0.019) or signal (*η*^2^ = 0.011) on the same metric. Signed hybrid achieves the highest mean bridge hub rank (0.964), followed by unsigned (0.929) and signed (0.871). The mechanism is the soft-power selection: signed requires high powers (mean 9–10) that suppress moderate positive correlations, which is precisely the correlation profile of bridge genes driven by a diluted mixed eigengene; signed hybrid hard-zeros negative correlations and achieves scale-free fit at lower powers (mean 2–7), preserving bridge gene cross-species adjacency weights. Despite this hub rank advantage, signed hybrid’s co-separation conditional on successful runs (0.66–0.71) is comparable to unsigned (0.78) and signed (0.70), because co-separation depends on the stability of the module community around bridge genes rather than bridge gene rank alone.

### 3.10 Library ratio imposes asymmetric species-level degradation

Within the bridge gene class, pathogen-side bridge genes consistently achieve higher *k*_cross_ rank than host-side bridge genes across all library ratios (Fig. 9). At high signal, *Blumeria* bridge genes achieve mean rank 0.985–0.994 while barley bridge genes achieve 0.817–0.912. This asymmetry is a read-depth artefact: cross-species connectivity for a *Blumeria* gene is the sum of its adjacency weights to barley genes, which remain at high read depth regardless of the ratio. As the ratio increases, within-species *Blumeria* correlations compress, narrowing the competition for the top of the *k-*_cross_ ranking. Barley and *Blumeria* bridge genes show nearly identical detection rates (81% at ratio 1:1, 80% at ratio 20:1 for both species at high signal, Fig. 8), confirming that collapse—not read depth asymmetry—is the primary driver of grey assignment. Consequently, in real high-ratio datasets, pathogen genes will systematically appear as stronger cross-species hubs than host genes, not because the biology is asymmetric but because the ranking reflects read-depth composition.

**Figure 7:**
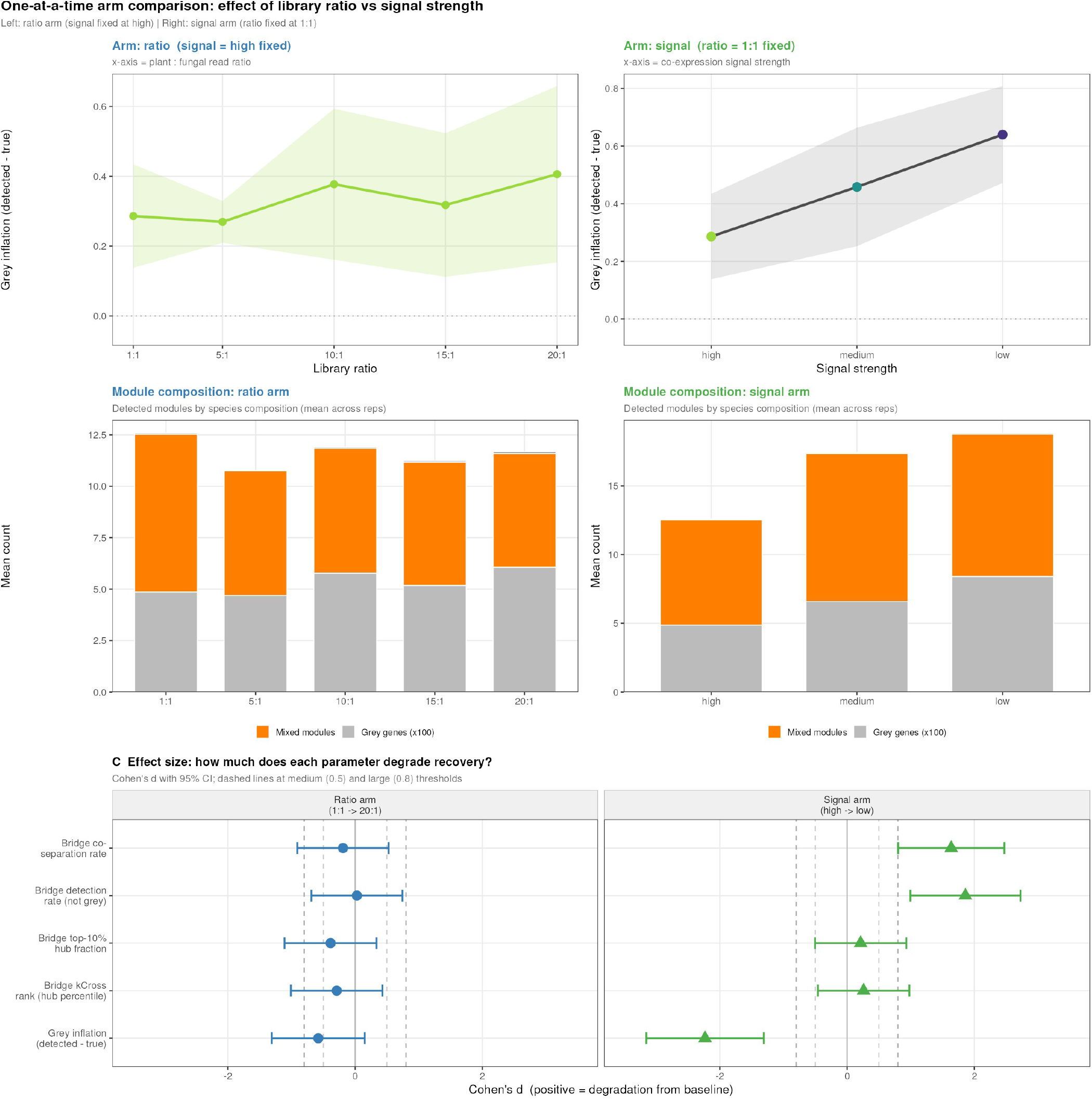
One-at-a-time arm analysis. **(Top row)** Grey inflation vs. ratio (left, signal fixed at high) and vs. signal level (right, ratio fixed at 1:1). All three network types produce overlapping trajectories. **(Middle row)** Per-run module composition as stacked bars (green = barley, purple = *Blumeria*, orange = bridge). Mixed-species modules dominate in both arms regardless of conditions. **(Bottom row)** Cohen’s *d* effect sizes comparing extreme vs. baseline parameter values for each bridge recovery metric. Ratio and signal show large effects on co-separation and detection rate; network-type differences are negligible.

**Figure 8:**
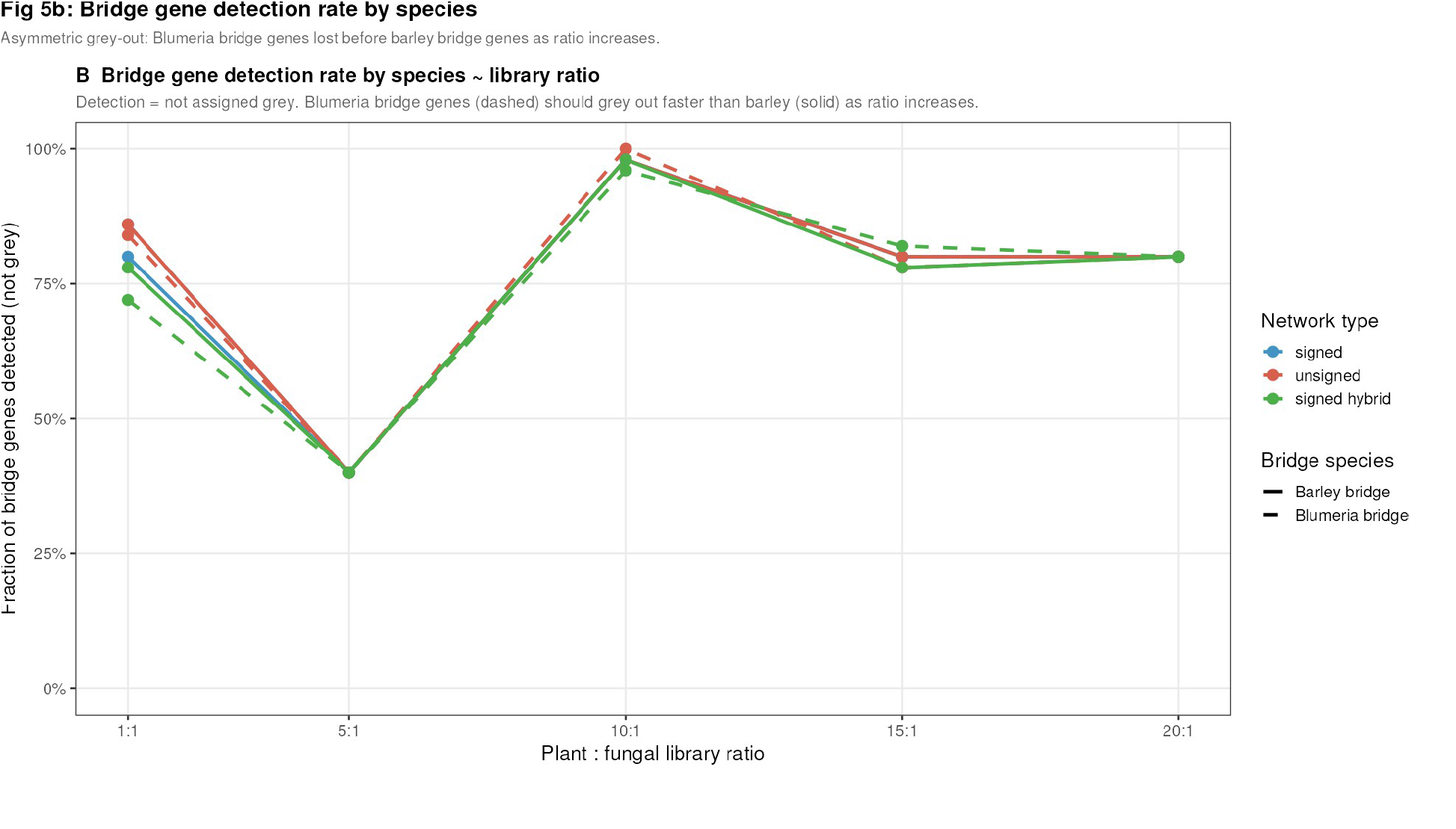
Bridge gene detection rate by species. Fraction of bridge genes not assigned to grey, stratified by species (barley solid, *Blumeria* dashed) and network type (colour), for the ratio arm (signal = high fixed). Detection rates for barley and *Blumeria* bridge genes are similar across the ratio range; collapse rather than read-depth asymmetry drives grey assignment.

**Figure 9:**
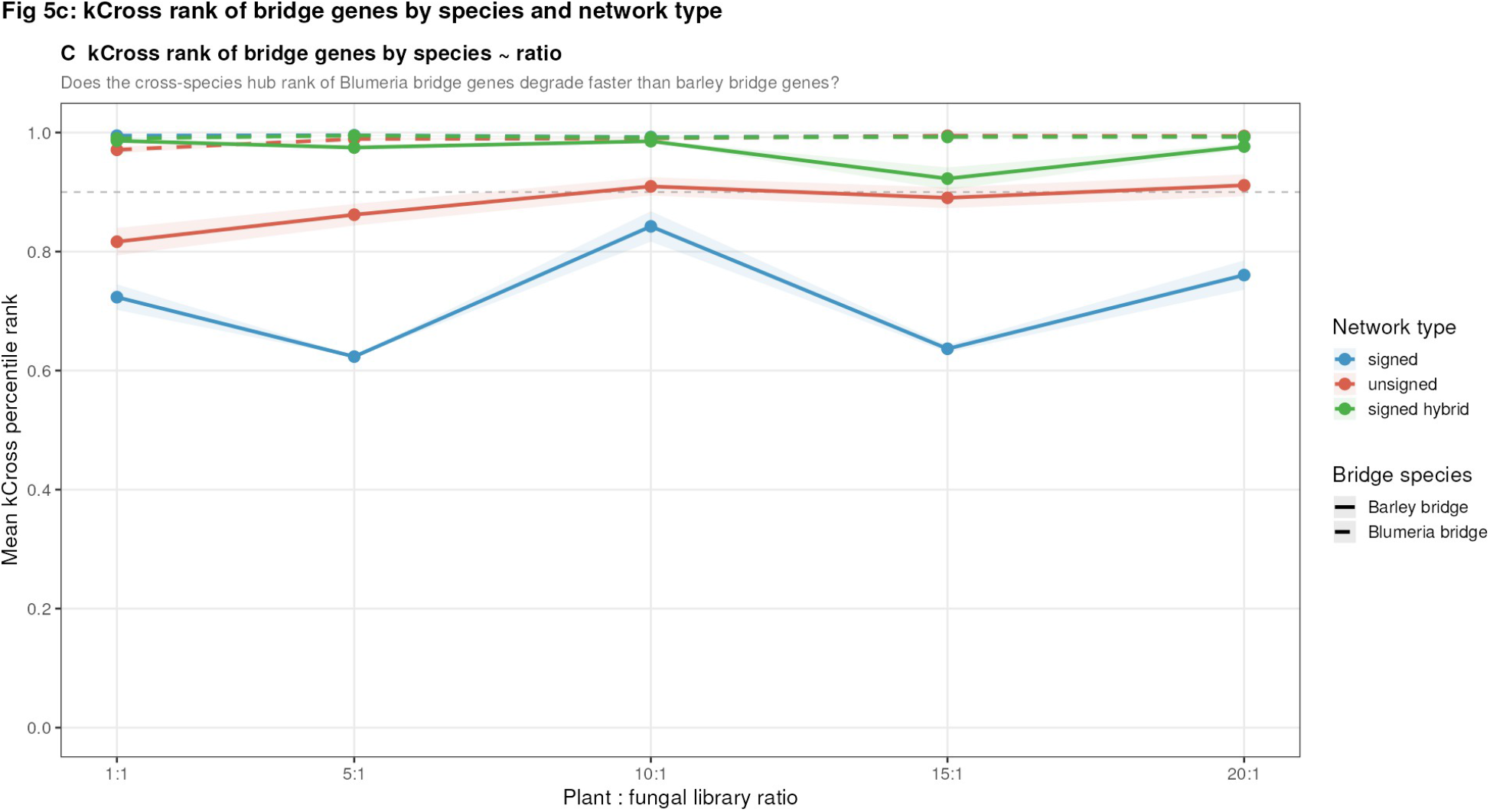
Cross-species hub rank of bridge genes by species. Mean *k*_cross_ percentile rank of bridge genes stratified by species, shown across library ratios for each network type (signal = high fixed). *Blumeria* bridge genes (dashed) achieve substantially higher rank than barley bridge genes (solid) at all ratios, reflecting a read-depth bias: *Blumeria* bridge genes are ranked by their connections to the deep barley side of the matrix.

### 3.11 Cross-species vs total connectivity identifies bridge genes as a distinct population

Figure 10 shows the relationship between total connectivity (*k*_total_) and cross-species connectivity (*k*_cross_) on a log–log scale for bridge, module, and grey genes at representative conditions underthe signed-hybrid approach. Bridge genes form a distinct cloud elevated above the module and grey gene distributions on the *k*_cross_ axis for a given *k*_total_, confirming that bridge genes are not merely highly connected in general but are specifically enriched for cross-species adjacency weight, consistent with their mixed-eigengene construction. This separation persists at ratios 5:1 and 10:1 but narrows at 20:1 as collapse events become more frequent.

**Figure 10:**
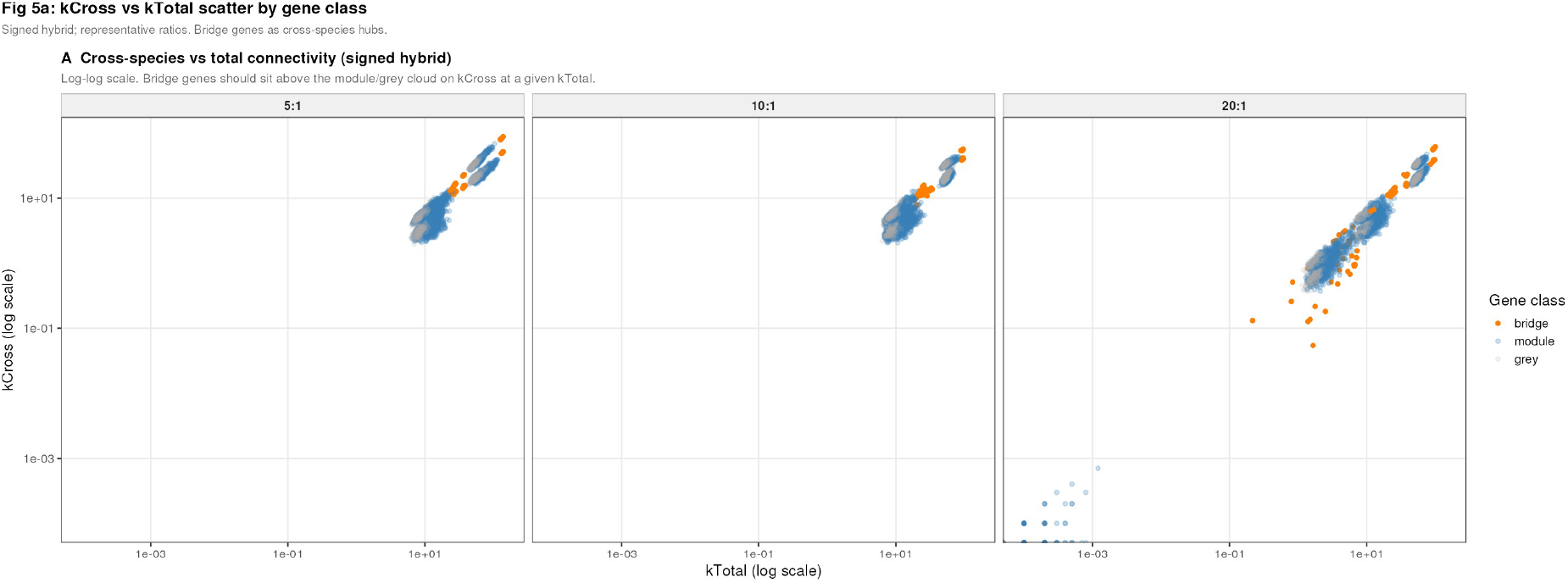
Cross-species vs. total connectivity by gene class. Log–log scatter of *k*_cross_ vs. *k*_total_ for bridge (orange), module (blue), and grey (grey) genes under signed hybrid at four representative library ratios (signal = high). Bridge genes form a consistently elevated cloud on *k*_cross_, confirming they are specifically enriched as cross-species hubs rather than generally highly connected genes.

### 3.12 Species composition of detected modules shifts with library ratio

In every condition where WGCNA detects non-grey modules, those modules contain genes from both species simultaneously — cross-species module membership is the default output of the method, not an exception. Figure~S1 shows how the minority-species fraction per module changes across the ratio and signal arms. At a balanced 1:1 ratio, the minority-species fraction within detected modules approaches 0.5, reflecting roughly equal plant and fungal representation. As the library ratio increases, the fungal fraction within modules declines systematically: at 20:1, detected modules still contain genes from both species, but fungal genes constitute a progressively smaller share of each module’s membership. This shift is a module-level manifestation of the same read-depth asymmetry that produces the hub rank bias described in Section~3.7 — as fungal read depth falls, fungal genes contribute less adjacency weight to the merged matrix, and their representation within co-expression communities is proportionally reduced even when they are not fully displaced to grey. The signal arm (right panel) shows that reducing co-expression signal strength shrinks the minority-species fraction further, compounding the ratio effect. Together these results confirm that library ratio and signal strength do not merely determine whether modules form, but actively shape the species composition of the modules that do form — a consideration that should inform how co-assigned effector--target pairs are interpreted in high-ratio experiments.

## 4 Discussion

We set out to evaluate whether single-network WGCNA, applied to a merged dual-organism expression matrix, can serve as a practical first-pass strategy for effector protein discovery — a novel application of the method for which no prior benchmark existed. The results support this use. Bridge genes, model effectors driven by a mixed host--pathogen eigengene, consistently rank in the 92nd percentile of cross-species connectivity across all tested conditions, and co-assign with their host interaction partners at a rate of 0.72 under favourable experimental conditions. The second contribution is a precise characterisation of when and why co-assignment succeeds or fails, providing actionable guidance for experimental design. The experiments motivating this work—finding effector proteins and characterising host–pathogen interaction interfaces—require the detection of *cross-species* co-expression, not separation of species. Reorienting the simulation around bridge genes—model effectors driven by a mixed eigengene with equal loading on host and pathogen expression programmes—reveals a more nuanced picture: the method can identify interaction-interface genes as connectivity hubs under a broad range of conditions, and co-assignment of effector–target pairs into shared modules succeeds in the majority of runs when signal quality is adequate — providing the first quantitative characterisation of the conditions under which dual-organism co-expression analysis is productive.

### Bridge genes are recoverable as cross-species hubs, but co-assignment fails under imbalance

The most important finding is that bridge genes are robustly identifiable as cross-species connectivity hubs regardless of library ratio or network type. Across all 225 simulation runs, bridge genes achieve a mean cross-species connectivity percentile rank of 0.92, substantially above module genes (0.50) and grey genes (0.45), and this separation is robust across all library ratios tested (Fig. 2). The hub signal persists because bridge genes carry strong co-expression with both the host and pathogen sides of the merged matrix via their mixed eigengene, and the soft-thresholded adjacency matrix preserves this structure even when fungal read depth is low. Critically, however, a high hub rank does not guarantee module co-assignment. Bridge co-separation falls from 0.73 at a balanced ratio and high signal to 0.32 at a 20:1 ratio (Fig. 5), and approaches zero under low-signal conditions. The distinction between being identifiable as a hub and being recoverable in a shared module reflects a fundamental compositionality limit: the adjacency matrix encodes the interaction signal, but module detection requires a coherent topological community on both sides of the interaction, and that community collapses under read depth asymmetry.

### Signal strength is the primary biological determinant of co-assignment success

The variance decomposition demonstrates that signal strength is the strongest driver of bridge co-separation (*η*^2^ = 0.12) and bridge detection (*η*^2^ = 0.07), and its effect operates primarily through the collapse mechanism: at low signal, between 47% and 93% of runs produce no usable module structure depending on the ratio. In collapsed runs, bridge hub rank is paradoxically *higher* (mean 0.96) than in successful runs (0.89), because the adjacency matrix is compressed into a single near-clique and bridge genes—which have genuine cross-species correlation—dominate the residual structure. This creates a practical trap: a collapsed run produces a hub ranking that looks informative but corresponds to no recoverable module biology. The ratio × signal interaction (*η*^2^ = 0.10 for co-separation) confirms that library imbalance only compounds an already-failed analysis; under low signal, increasing the ratio has no further negative effect because the method has already failed. Signal strength in real experiments reflects the quality and timing of infection samples, the degree of biological replication, and the dynamic range of the transcriptional response. Experimental design choices that improve these — synchronised inoculation, dynamic timepoint sampling, adequate replication — increase the probability that an experiment falls in the regime of reliable co-assignment, but cannot guarantee it: the underlying transcriptional signal is a property of the biology, not the experimenter.

### Library ratio imposes asymmetric degradation at the gene level

Among bridge genes, pathogen-side genes consistently achieve a higher cross-species connectivity rank (mean 0.98) than host-side bridge genes (mean 0.86) across all library ratios (Fig. 9). This counter-intuitive result arises because cross-species connectivity is measured as the sum of adjacency weights to the *other* species: a pathogen bridge gene’s cross-species connectivity is its summed weight to host genes, which remain at high read depth regardless of ratio. As the ratio rises, the pathogen module genes lose read depth and their within-species correlations collapse, pushing pathogen bridge genes to the top of the distribution. In real high-ratio datasets, pathogen genes will therefore systematically appear as stronger cross-species hubs than host genes due to read depth composition, not biology. Any candidate effector list derived from hub ranking in high-ratio data should be interpreted with this bias in mind.

### Network type affects hub detectability but not co-assignment

Network type accounts for 18% of variance in bridge hub rank—more than ratio or signal—but only 1% of variance in bridge co-separation (*η*^2^ = 0.011; Fig. 6C). Signed hybrid networks achieve the highest mean hub rank (0.964) because its low soft-threshold power (mean 2–7) preserves moderate correlations that bridge genes produce via their diluted mixed eigengene loading. Signed networks require much higher powers (mean 9–10), which suppress moderate correlations and reduce bridge gene ranks. Despite this advantage, signed hybrid’s bridge co-separation conditional on successful runs (0.66–0.71) is comparable to unsigned (0.78) and signed (0.70), because co-separation depends on the stability of the surrounding module community rather than bridge gene rank alone. The practical recommendation is to use signed hybrid as the default network type for dual-organism data—it offers the best chance of identifying effector candidates as hubs—while recognising that no network type circumvents the underlying community structure problem.

### Grey inflation identifies a specific target for methodological improvement

Even in successful runs, grey inflation is the most direct predictor of bridge co-separation failure (Pearson *r* = −0.35 across successful runs). Bridge genes are disproportionately susceptible because their mixed eigengene loading produces moderate rather than strong correlations with any single gene set—precisely the profile that the scale-free topology constraint and high soft powers target for suppression. Grey inflation is not a parameter tuning problem: it reflects the inherent competition between within-species correlation structure and the merged-matrix scale-free topology criterion, and no standard WGCNA parameter set resolves this competition in favour of cross-species bridge genes.

### Implications for published dual-organism co-expression analyses

Studies reporting plant–pathogen co-expression modules from jointly analysed RNA-seq data should be interpreted with awareness of three structural limits identified here: (1) hub rank in the adjacency matrix is a more reliable indicator of interaction-interface genes than module co-assignment, which is sensitive to collapse; (2) pathogen genes will be systematically ranked higher as cross-species hubs in high host-to-pathogen ratio datasets due to read depth asymmetry, not biology; and (3)the strength of the underlying co-expression signal, which experimental design can improve but not fully determine, is the primary biological factor governing whether the analysis is interpretable.

### Towards interaction-aware co-expression analysis

The simulation framework developed here provides a ready-made benchmark for evaluating species-aware methods. Two strategies address different aspects of the problem. Running WGCNA separately on each species’ expression matrix (normalised independently) avoids signal mixing entirely and is appropriate when within-species module structure is the primary research question. For cross-species interaction discovery, structured methods that explicitly model species membership—sparse canonical correlation, cross-species biclustering, or multi-view clustering—could recover interaction modules without conflating them with intraspecies structure. Any proposed method should be evaluated against the bridge co-separation metric introduced here, which directly measures recovery of the interaction interface. The asymmetric degradation finding further suggests that methods should explicitly account for read depth imbalance in their weighting of cross-species correlations.

### Limitations

Our simulation uses a fixed dispersion (*α* = 0.1) and AR(1) autocorrelation (*ϕ* = 0.7) that may not capture the full variance of infection time courses. The gene pool is small relative to a full transcriptome, the number of bridge genes is minimal (20 total), and the bridge eigengene uses symmetric 0.5:0.5 loading that may not reflect the asymmetric effector biology of real host–pathogen interactions. We do not model batch effects, zero-inflation, or within-replicate infection heterogeneity. Future work should extend the simulation to realistic transcriptome scales, asymmetric bridge eigengene architectures, larger bridge gene sets, and alternative network construction methods including partial correlation and distance-based networks. The bridge co-separation metric should also be validated against real dual-organism datasets where effector–target gene pairs are independently known.

## 5 Conclusions

We present a simulation benchmark for dual-organism co-expression analysis that but their co-assignment to shared detected modules is fragile and fails in approximately 41% of runs. The central finding is that WGCNA can identify effector-like genes as cross-species connectivity hubs under a broad range of conditions—bridge genes achieve mean hub rank in the 92nd percentile across all 225 simulation runs—but their co-assignment to shared detected modules is fragile and fails in approximately 41% of runs due to scale-free topology when the network was simulated with a low signal.. Co-separation, the fraction of host–pathogen bridge gene pairs assigned to the same detected module, is the most biologically meaningful metric and the one most sensitive to biological and technical conditions: signal strength (*η*^2^ = 0.12) and library ratio (*η*^2^ = 0.22) are its primary determinants, with network type contributing less than 2%.

Three compositionality limits are responsible for these failures. First, the soft-power selection operates on a single merged matrix and cannot simultaneously satisfy the scale-free topology criterion for both species’ correlation landscapes, causing approximately 41% of runs to collapse regardless of network type or signal level. Second, library ratio asymmetrically degrades the pathogen side of the bridge co-expression signal, leading to preferential grey assignment and loss of the topological community needed for co-assignment—a failure distinct from hub rank loss, which is preserved due to relative ranking effects. Third, grey inflation selectively suppresses moderate-correlation genes—the profile shared by bridge genes—even in successful runs, reducing co-separation by approximately 3.5 percentage points per 10% increase in grey inflation.

Above all, experimental design—ensuring adequate pathogen read depth and selecting timepoints that capture dynamic transcriptional responses—is more effective than any analytical choice. Signed hybrid is the preferred option for hub detection (highest bridge hub rank, *η*^2^ = 0.18 for network type on rank), but its advantage does not extend to co-assignment. The practical recommendation for effector discovery studies is to use signed hybrid for hub ranking, to interpret high cross-species connectivity rank as the primary evidence for interaction-interface candidacy, and to treat module co-assignment as confirmatory rather than primary evidence. Above all, experimental design—ensuring adequate pathogen read depth and selecting timepoints that capture dynamic transcriptional responses—is more effective than any analytical choice in determining whether interaction-interface genes can be recovered.

## 6 Methods

### 6.1 Simulation framework overview

We designed a simulation pipeline to generate dual-organism RNA-seq count matrices with known ground-truth co-expression module structure, targeted at the biological objective of recovering effector-like genes at the host–pathogen interaction interface. The pipeline proceeds in six stages:

1. sampling real gene models from reference annotations,
2. simulating module eigengene time courses using autoregressive AR(1) processes with replicate-level noise,
3. constructing gene-level expected expression matrices with hub-to-peripheral correlation gradients,
4. generating RNA-seq counts using the `polyester` simulation framework [Frazee et al., 2015] with realistic negative-binomial overdispersion and library size imbalance,
5. applying species-separated variance-stabilising transformation (VST) normalisation via DESeq2 [Love et al., 2014], and
6. running WGCNA with three network construction types on the merged normalised matrix. All analyses were performed in R 4.3.2 using WGCNA 1.74 [Langfelder and Horvath, 2008], DESeq2 1.42 [Love et al., 2014], polyester 1.38 [Frazee et al., 2015], aricode 1.0, and parallel 4.3.2 for multi-core execution. Scripts and simulation outputs are available at [repository URL].

### 6.2 Gene model sampling

Gene lengths and identifiers were drawn based on the annotated gene models of both species: the *H. vulgare* MorexV3 (Mascher et al. 2021) (downloaded from Ensembl Plants release 62) and the *B. graminis* f.sp. *hordei* DH14v4 annotation (REF)For each organism, gene length was defined as the maximum exon-to-exon span per gene identifier. A fixed random sample of 625 barley genes (5 modules × 100 genes/module + 125 grey genes) and 375 *Blumeria* genes (3 modules × 100 genes/module + 75 grey genes) was drawn without replacement (seed 42) and held constant across all simulation conditions, giving a total pool of 1,000 genes. An additional 20 *bridge genes* (10 barley + 10 *Blumeria*) were designated as model effectors; their expression was constructed from a mixed eigengene with equal loading on the first eigengenes of both species (see Section 2.3). Bridge genes were drawn from the existing module gene pool and overwritten during expression construction, so the final gene pool comprises 500 barley module genes, 125 barley grey genes, 300 *Blumeria* module genes, and 75 *Blumeria* grey genes, with 20 of the module genes reclassified as bridge class at the point of overwrite. Each gene was assigned one of three ground-truth classes: *bridge* (overwritten with mixed eigengene), *module* (pure single-species co-expression structure), or *grey* (background, no module structure). Grey fraction was fixed at 20% per species.

### 6.3 Eigengene simulation with replicate structure

Module eigengene trajectories were simulated as autoregressive AR(1) processes at the timepoint level, with biological replicate noise added within each timepoint. The simulation used *T* = 5 timepoints and *R* = 12 replicates per timepoint, yielding *n* = 60 samples. For each module *m*, the timepoint-level trajectory was:

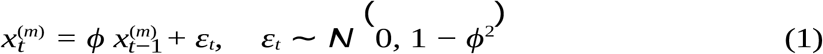

with *ϕ* = 0.7 (moderate temporal autocorrelation). Each sample-level expression value was then:

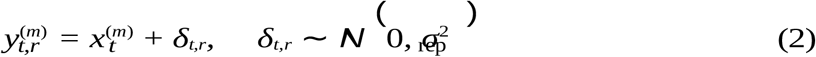

with *σ*_rep_ = 0.3. Each eigengene was *z*-scored to zero mean and unit variance after replicate expansion. This design produces a block-structured correlation matrix that more closely resembles a real factorial RNA-seq infection time course than a simple continuous-trajectory simulation, and a fresh set of eigengenes was generated per simulation replicate using an independent random seed.

Bridge genes received a mixed eigengene constructed as an equal-weighted sum of the first plant and first fungal eigengenes, normalised to unit variance:

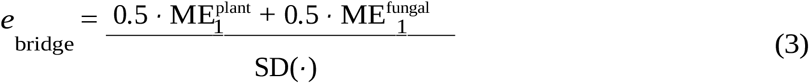

This gives bridge genes moderate co-expression with both species simultaneously, analogous to a secreted effector that is transcriptionally co-regulated with the infection programme on both sides.

### 6.4 Gene-level expression with modular correlation structure

For gene *g* assigned to module *m*, the expected expression was:

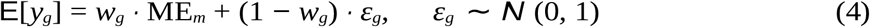

where *w*_*g*_ is the gene’s module membership weight, drawn uniformly from [*ρ*_min_, *ρ*_max_] per signal level. Three signal levels were defined:

- **High signal**: *ρ* ∈ [0.50, 0.90]
- **Medium signal**: *ρ* ∈ [0.30, 0.70]
- **Low signal**: *ρ* ∈ [0.10, 0.50]

This produces realistic within-module correlation gradients from hub to peripheral gene. Grey genes were assigned *w*_*g*_ = 0 and drawn from independent Gaussian noise. Expected expression vectors were scaled to per-species target library totals using gene length and library size as described in Section 2.5.

### 6.5 RNA-seq count simulation with library size imbalance

RNA-seq counts were generated using the `polyester` package [Frazee et al., 2015], which simulates negative-binomial counts with length bias. Library sizes were drawn from *N* (10 × 10^6^, (1 × 10^6^)^2^) per sample, clipped to a minimum of 5 × 10^6^. Count matrices for barley and *Blumeria hordeii* were generated independently using real gene lengths from the reference annotations. Library size imbalance was imposed by partitioning total reads between species:

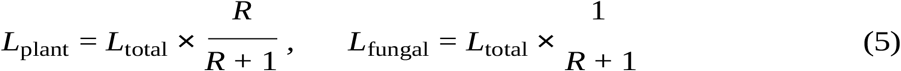

where *R* is the plant-to-fungal read ratio. Overdispersion was modelled via `polyester`’s dispersion function with a fixed log-intercept of log(0.1) (corresponding to negative-binomial dispersion parameter *α* = 0.1); on rare numerical failures the fallback `rnbinom(size = 10)` was used. Counts from both species were kept as separate matrices and normalised independently before merging for WGCNA input.

### 6.6 Normalisation and preprocessing

Each species’ count matrix was normalised independently using DESeq2’s variance-stabilising transformation with no experimental design (`design = ~1`, `blind = TRUE`) [Love et al., 2014]. Size factors were estimated via the median-of-ratios method; genes with all-zero counts were removed before VST. The two species-specific VST matrices were then merged by matching shared sample identifiers, transposed to samples-by-genes orientation (60 × 1,000), and passed directly to WGCNA. Species-separated normalisation prevents host read depth from dominating the size factor estimates for the minority-species matrix.

### 6.7 WGCNA network construction and module detection

For each merged normalised matrix, WGCNA was applied under all three standard network construction types: **signed, unsigned**, and **signed hybrid**. The transformations from biweight midcorrelation *r* to soft-thresholded adjacency *a* are:

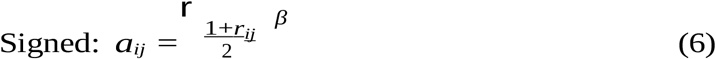

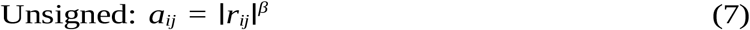

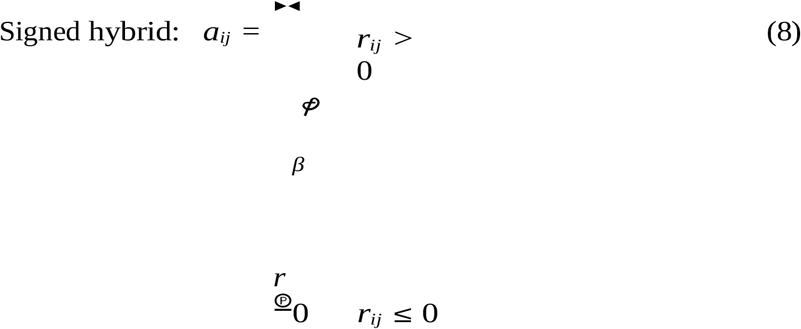

All three methods used biweight midcorrelation (`bicor`, `maxPOutliers = 0.1`, pairwise-complete observations). The TOM type was held fixed at “`signed`” across all three network types to isolate the effect of the adjacency transformation. Soft-thresholding power *β* was selected independently per run and per network type via `pickSoftThreshold` over the range *β* ∈ [1, 30], targeting scale-free fit *R*^2^ ≥ 0.80; when no power achieved this threshold, the power with the highest *R*^2^ was used. Modules were detected via `blockwiseModules` with `minModuleSize = 15`, `mergeCutHeight = 0.35`, `pamRespectsDendro = FALSE`, and `numericLabels = FALSE`. Each WGCNA call used `nThreads = 1` to avoid oversubscription within the parallel worker pool. Critically, each simulated dataset was analysed by all three network types from the same VST matrix, with a fresh `pickSoftThreshold` call per type, ensuring that differences reflect the adjacency transformation and power selection rather than differences in the underlying data.

### 6.8 Experimental design

The simulation tested a four-arm factorial design to isolate and cross the effects of library ratio and co-expression signal strength (Fig. 1).

#### Baseline arm

(`ratio = 1:1`, `signal = high`): a single condition establishing the performance ceiling under ideal conditions.

#### Ratio arm

(`signal = high` fixed, `ratio` ∈ *{*1, 5, 10, 15, 20*}*): isolates the pure effect of host:pathogen read-depth imbalance at best-case signal strength.

#### Signal arm

(`ratio = 1:1` fixed, `signal` ∈ *{*high, medium, low*}*): isolates the pure effect of co-expression signal strength at balanced read depth.

#### Full factorial arm

(all 15 ratio × signal combinations): enables two-way variance decomposition and interaction detection, and subsumes the three arms above as special cases.

The baseline condition (`ratio = 1`, `signal = high`) appears in all four arms and serves as the shared anchor for cross-arm comparisons. After deduplication, the four arms yielded 15 unique (ratio, signal) combinations, each replicated 5 times with independent random seeds and analysed under all three network types, producing 15 × 5 × 3 = 225 total WGCNA calls from 75 independently simulated expression datasets. Simulations were executed in parallel across 6 CPU cores via `parallel::mclapply`, with WGCNA’s internal thread pool disabled.

### 6.9 Hub connectivity metrics

In addition to module-level assignment metrics, per-gene connectivity was computed from the soft-thresholded adjacency matrix using the same `bicor`-based adjacency function and the network-type-specific *β* already selected for that run. Self-loops were zeroed before computing connectivity. For gene *g*:

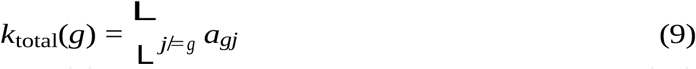

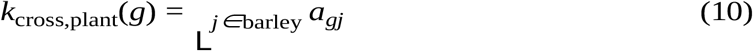

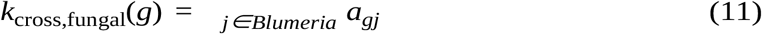

Cross-species connectivity was defined directionally: for a barley gene, *k*_cross_(*g*) = *k*_cross,fungal_(*g*); for a *Blumeria* gene, *k*_cross_(*g*) = *k*_cross,plant_(*g*). Percentile ranks 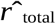 and 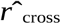 were computed within each run (average-method ranking divided by the total number of genes). These pergene statistics are reported in Table SX for all five replicates and support the species-stratified bridge rank analyses in Figs. 3–8.

### 6.10 Performance metrics

Module recovery and bridge gene detectability were quantified using the following metrics.

#### Adjusted Rand Index (ARI)

Overall agreement between true and detected module labels, corrected for chance, computed via `aricode::ARI`. Retained as a global partition quality diagnostic.

#### Grey inflation

Difference between the detected and ground-truth grey fractions 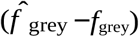. Positive values indicate excess grey assignment. Visualised in Figs. 7 and 6.

#### Species contamination

Fraction of detected non-grey modules containing genes from both species simultaneously. Retained as a diagnostic for module composition visualisations (Fig. 7B); not used as a primary evaluation metric because cross-species modules represent the target, not a defect.

#### Bridge cross-species hub rank

(`bridge kCross rank`): Mean percentile rank of all 20 bridge genes in the within-run *k*_cross_ distribution. The null expectation is 0.50; values near 1 indicate bridge genes rank as the strongest cross-species hubs (Fig. 2).

#### Bridge top-10% hub fraction

(`bridge top10 frac`): Fraction of bridge genes in the 90th percentile or above of the *k*_cross_ distribution, with null expectation 0.10 (Fig. 4).

#### Bridge detection rate

(`bridge detected frac`): Fraction of bridge genes not assigned to the grey module. A bridge gene assigned to grey cannot co-cluster with its interaction partner, so this is a prerequisite for co-separation (Fig. 8).

#### Bridge co-separation

(`bridge coseparation`): Fraction of all possible cross-species bridge gene pairs (10 plant × 10 fungal = 100 pairs) in which both members are assigned to the same detected non-grey module. This is the primary evaluation metric, directly testing whether the method co-assigns model effectors and their host interaction partners (Figs. 5 and 6).

### 6.11 Variance decomposition

To quantify the relative contributions of library ratio, signal strength, and network type to each performance metric, we fitted a fully saturated three-factor linear model to all 225 simulation runs:

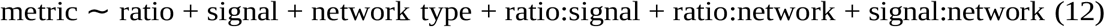

Eta-squared (*η*^2^ = *SS*_term_*/SS*_total_) was computed for each term from the sequential ANOVA decomposition. The residual *η*^2^ reflects replicate-to-replicate variability within conditions (Table Sx, Fig. 6C).

### 6.12 Output checkpoint and reproducibility

Before executing any simulation runs, the script verifies whether a complete, internally consistent result set already exists in `results/wgcna benchmark v7/`. Five conditions are checked: all five output files are present; `benchmark results v7.csv` contains exactly 225 rows; all condition × replicate × network-type combinations are represented; all required columns are present; and no NA values appear in the four primary bridge recovery metrics. If all checks pass, the simulation is skipped and results are loaded from disk; if any check fails, the specific reason is printed and the full simulation is re-run from scratch. Random seeds are derived deterministically from the simulation run index (`seed base` = `run i` × 10,000), guaranteeing reproducibility regardless of parallel scheduling order. The complete pipeline is implemented in scripts `03.1 dual species wgcna benchmark sim v7.R` (simulation) and `03.3 narrative figures v7.R` (figures).

## Data and code availability

Simulation code, annotation parsing scripts, visualisation scripts, and analysis outputs are available at https://github.com/afenn/dual-species-wgcna-benchmark (to be made public upon acceptance). Reference genome annotations are publicly available from Ensembl Plants (barley MorexV3, release 62) and the *B. graminis* genome database (https://blumeria.net). All simulations were run on the Leibniz Supercomputing Centre (LRZ) CoolMUC-4 cluster using 6 CPU cores per job. Raw simulation results (225 runs × per-run metrics) are provided as CSV files in the GitHub repository.

## Author contributions

A.F.: conceptualisation, methodology, software, formal analysis, visualisation, writing (original draft). N.K.: conceptualisation, supervision, writing (review and editing). R.H.

## Acknowledgements

We thank the Leibniz Supercomputing Centre (LRZ) for computational resources and Peter Langfelder for helpful discussions on module preservation statistics. A.F. is supported by the German Research Foundation (DFG) - TRR 356/1 2023 – 491090170

## Supplementary Figures

**Figure S1:**
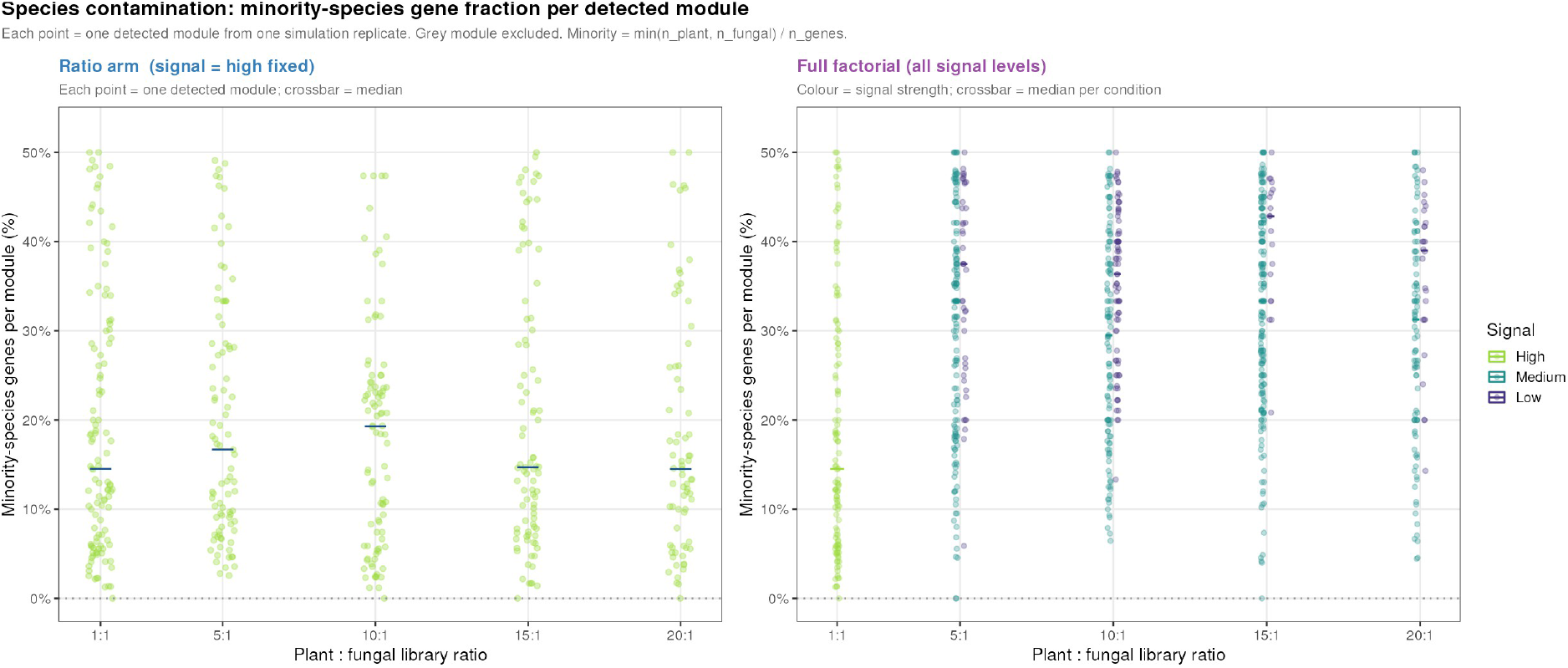
**S**pecies composition of detected modules across ratio and signal arms. Each point represents one detected non-grey module from one simulation replicate; crossbar = median per condition. Minority-species fraction is defined as min(nplant,nfungal)/ngenes per module. **(Left)** Ratio arm (signal = high fixed): minority-species fraction declines with increasing library ratio, reflecting the progressive underrepresentation of fungal genes within co-expression communities as read depth falls. **(Right)** Full factorial (all signal levels): signal strength compounds the ratio effect, with low signal producing modules with the smallest fungal representation. Cross-species module membership persists across all conditions, but its composition is systematically distorted by both library ratio and signal strength.

**Figure S2:**
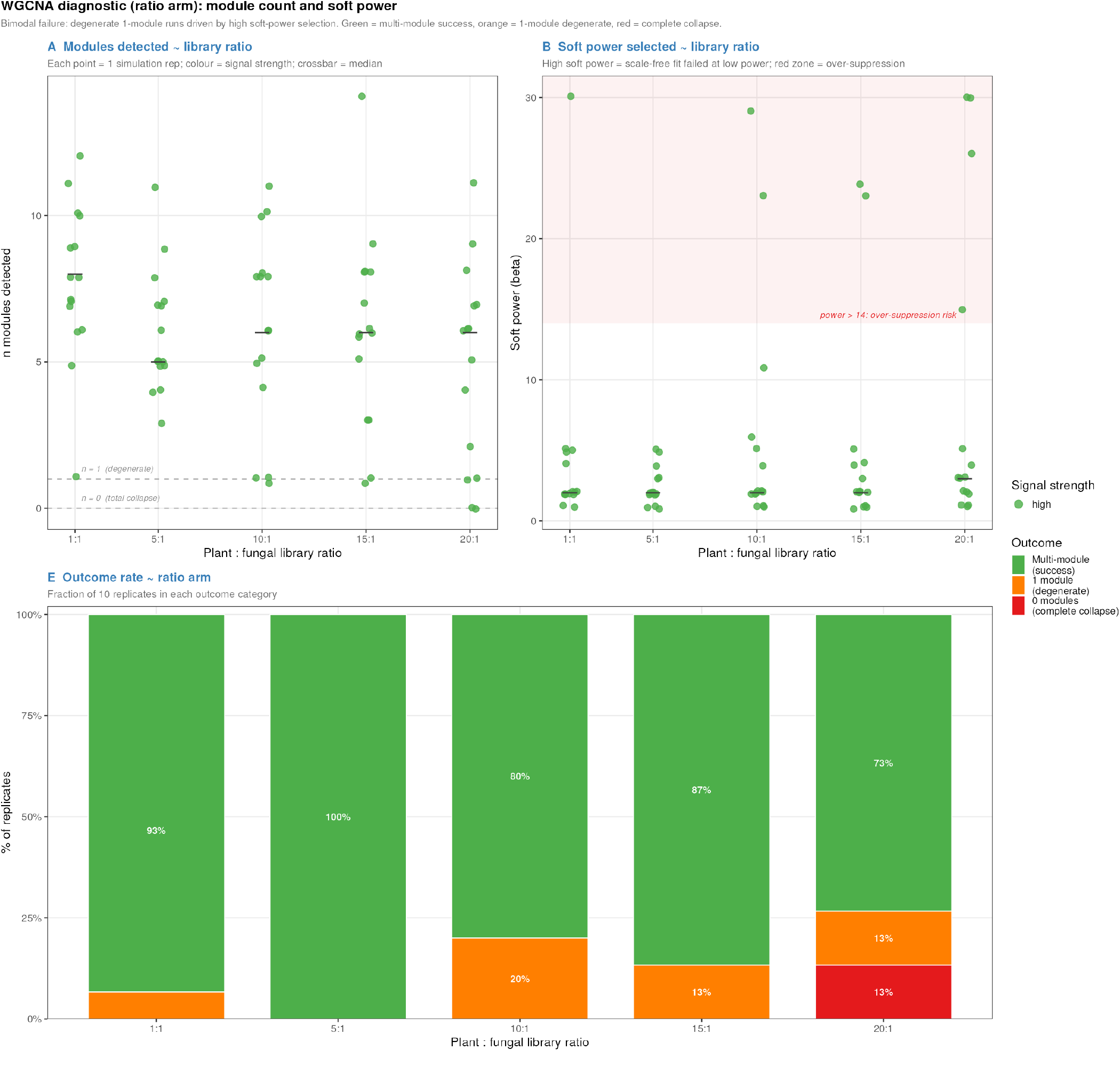
WGCNA diagnostic statistics across library ratio (ratio arm, signal = high fixed). **(A)** Number of detected modules per simulation replicate as a function of library ratio. Dashed lines mark *n* = 0 (total collapse) and *n* = 1 (degenerate single-module outcome). **(B)** Soft-thresholding power *β* selected by `pickSoftThreshold` per replicate. The red shaded zone (*β >* 14) indicates conditions where over-suppression of moderate correlations is likely. **(E)** Stacked bars showing the fraction of replicates in each outcome category (successful multi-module detection, degenerate, or collapsed) per ratio level. Each point = one simulation replicate; crossbar = median.

**Figure S3:**
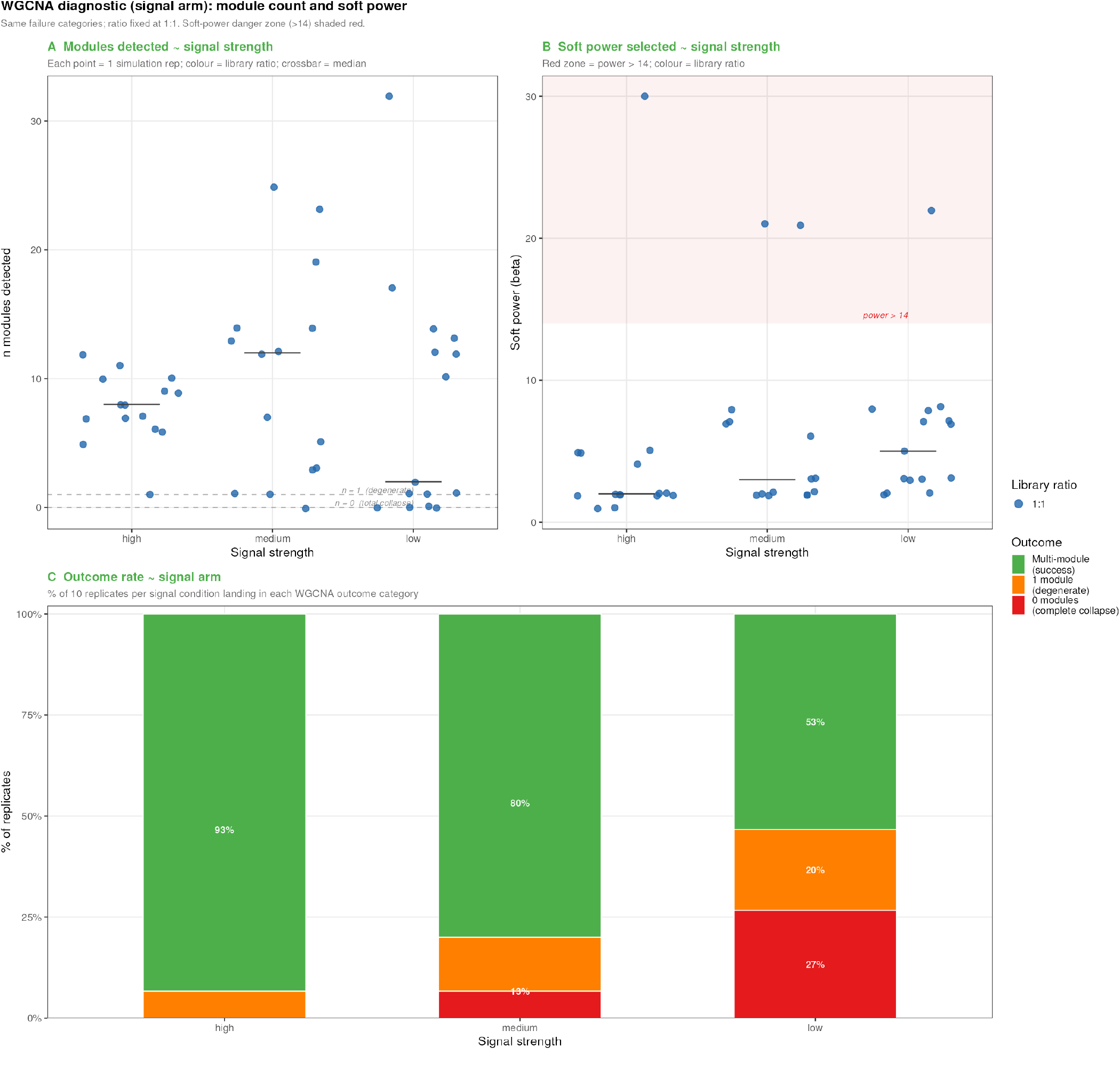
WGCNA diagnostic statistics across signal strength (signal arm, ratio = 1:1 fixed). **(A)** Number of detected modules per simulation replicate as a function of signal level. **(B)** Soft-thresholding power *β* selected per replicate; colour indicates library ratio. At low signal, soft powers frequently reach the ceiling of the search range (*β* = 30), consistent with scale-free topology never being achieved. Each point = one simulation replicate; crossbar = median.

